# DEET feet: *Aedes aegypti* mosquitoes use their tarsi to sense DEET on contact

**DOI:** 10.1101/360222

**Authors:** Emily Jane Dennis, Leslie Birgit Vosshall

## Abstract

DEET (*N*, *N*-diethyl-*meta*-toluamide) is the most effective and broadly used insect repellent, but its mechanism of action is both complex and controversial [1]. Previous work demonstrated that DEET acts both on insect smell [2-6] and taste [7-11] systems. Its olfactory mode of action requires the odorant co-receptor *orco* [2, 3, 6], while its gustatory repellency is mediated by activation of bitter taste receptors and neurons in the proboscis upon ingestion [8]. Together, these data have led to the assumption that DEET acts only on olfactory and gustatory pathways. We previously observed that *orco* mutant female *Aedes aegypti* mosquitoes are strongly attracted to humans even in the presence of DEET, but are rapidly repelled after contacting DEET-treated skin [6]. To understand the basis of this contact chemorepellency, we carried out a series of behavioral experiments and discovered that DEET acts in three distinct ways: through smell, taste, and contact. DEET and bitter tastants are feeding deterrents when ingested, but only DEET is capable of mediating contact repellency on human skin. We show that the repellent touch of DEET is mediated by the tarsal segments of the legs, and not gustatory neurons in the proboscis as previously believed. This work establishes mosquito leg appendages as the actual sensors of DEET, and highlights the existence of an unknown sensory pathway that is independent of bitter taste. These results will inform the search for novel contact-based insect repellents.

**Highlights:**

- DEET and bitters are both repellent when ingested by *Aedes aegypti* female mosquitoes
- Only DEET is additionally repellent upon contact
- Repellency of DEET on skin is mediated solely by the legs
- Any of the three pairs of legs can sense DEET and prevent mosquitoes from biting

## Results and Discussion

### DEET and Bitter Compounds are Repellent When Ingested, but only DEET is Repellent on Contact

*orco* mutant mosquitoes are repelled by DEET on contact [6] but the sensory appendages, sensory neurons, and chemosensory receptor genes required for this phenomenon are unknown. To study contact repellency, we used heteroallelic *orco^5/16^* mutant mosquitoes throughout this study to eliminate the olfactory effects of DEET. Here we define “olfactory” repellency” as avoidance of volatile DEET that is dependent on *orco*, “taste” and “gustatory repellency” as the anti-feedant effect seen after ingesting fluid, and “contact repellency” as the repellency of a surface, usually but not exclusively human skin.

Both *D. melanogaster* flies [8] and *Ae. aegypti* mosquitoes [12] will reject sucrose tainted with bitter substances. Previous work in *D. melanogaster* flies [8] demonstrated that DEET acts as a bitter tastant when ingested, and that bitter-sensitive taste neurons and gustatory receptor genes required to sense bitters are responsible. Similar bitter- and DEET-sensitive neurons were identified on the proboscis of the mosquito [10] but their influence on behavior is unknown. To ask if DEET can inhibit *Ae. aegypti* mosquito sugar feeding, we offered females a choice between drinking either untainted 10% sucrose or 10% sucrose mixed with 1% DEET or bitters (1 mM lobeline or 5 mM quinine) in a CAFE assay [12-14] (Fig. 1A). Fasted female mosquitoes avoided both bitter tastants and DEET (Fig. 1B-D). These data demonstrate that in mosquitoes, as in *D. melanogaster* flies [8] and *Apis mellifera* bees [7], DEET and bitter tastants induce avoidance of an otherwise attractive sucrose solution.

**Figure 1.**
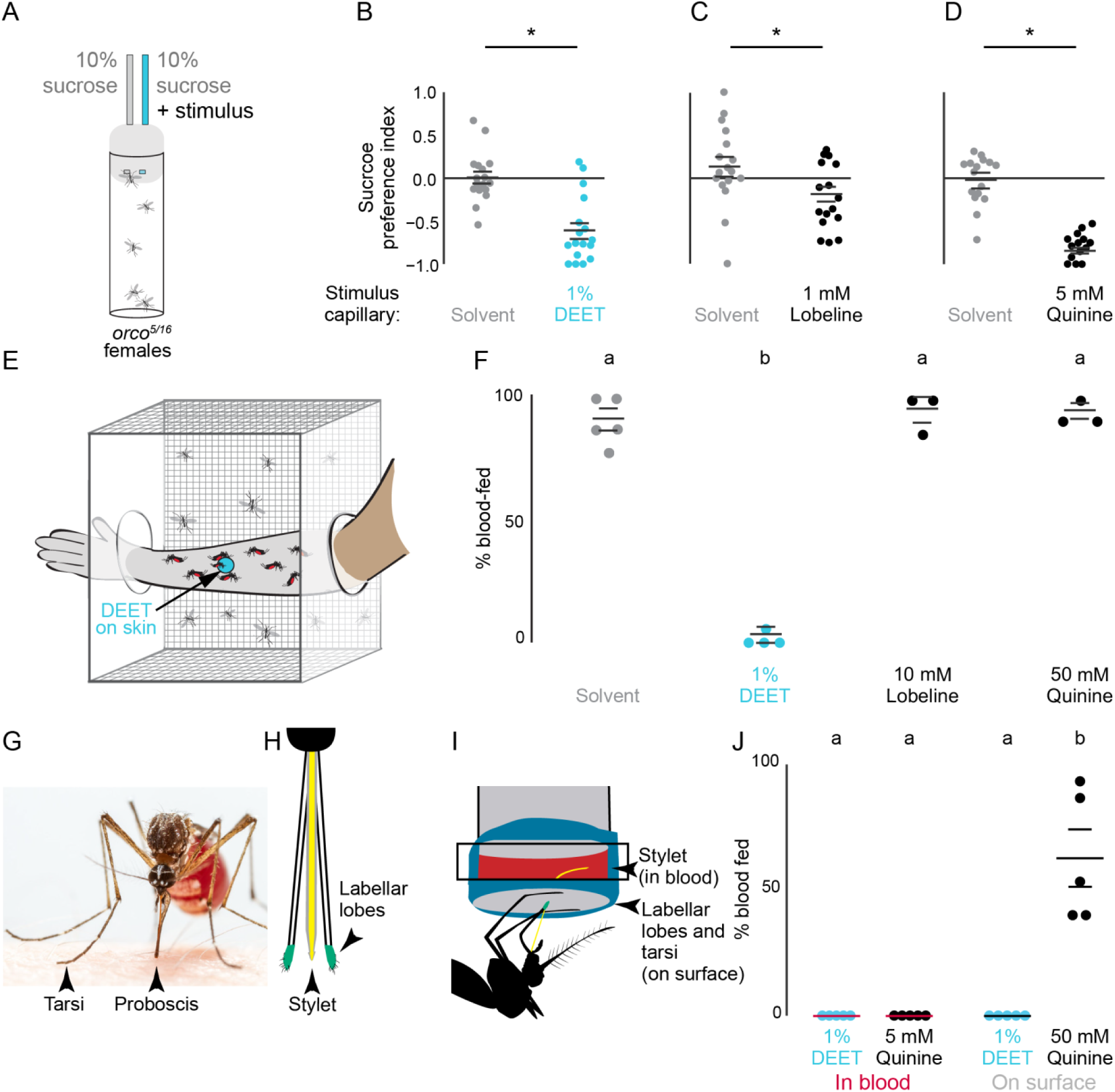
| DEET and Bitter Compounds are Repellent When Ingested, but only DEET is Repellent on Contact. (A) Mosquito CAFE assay schematic. (B-D) Inhibition of sucrose ingestion by DEET (B), lobeline (C), or quinine (D) in the CAFE assay with solvent controls indicated by the gray dots (N=14-17, n=5 females/assay). (E) Arm-in-cage schematic of a DEET-treated arm with a 25 mm circle of accessible skin. (F) Blood-feeding with the indicated compounds applied to a human arm as in (E) (N=3-5, n=23-25 females/assay). (G) Female mosquito feeding on a human arm with proboscis contacting the skin. (H) Schematic of the sensory appendages of the mosquito proboscis. (I) Glytube assay schematic highlighting location of appendages during feeding. (J) Glytube feeding with indicated compounds applied in blood (left) or on surface (right) (N=5, n=12-16 females/assay). Horizontal lines in B-D, and J represent mean ± SEM. Different letters or ^∗^ indicate statistically significantly distinguishable groups (p<0.05 Student’s *t*-test (B-D), or one-way (F) or two-way (J) ANOVA with Tukey’s post-hoc test.

To ask if bitter taste alone accounts for the contact chemorepellency of DEET on skin, we used a modified arm-in-cage assay [6, 15, 16] (Fig. 1E) in which mosquitoes were offered the opportunity to blood-feed on a 25 mm circle of exposed skin treated with solvent, DEET, lobeline, or quinine. Remarkably, applying either bitter tastant to skin had no effect on mosquito biting and blood-feeding behavior (Fig. 1F), even though they were delivered at 10-fold higher concentrations than those that deterred sugar feeding. In contrast, DEET applied on the arm provided complete protection (Fig. 1F). This is strong evidence that DEET is not activating bitter taste pathways to repel insects from skin. Instead, there must be an unknown third sensory system, apart from smell and bitter taste that mediates contact chemorepellency.

To reconcile how bitters can be effective anti-feedants in sugar-feeding assays but not in blood-feeding assays, we tested if the delivery of the bitter tastant is the salient difference between the two assays. Mosquitoes contact skin surface with their legs and proboscis (Fig. 1G). To bite a human arm, a mosquito must first saw through the skin and insert a needle-like appendage, the stylet, under the skin (Fig. 1H). Therefore, only the stylet is able to contact the blood. We hypothesized that bitter tastants may be effective on taste, here defined as ingestion through the stylet, but not on contact.

To test this hypothesis, we used a Glytube feeding assay (Fig. 1I). The Glytube assay uses a piece of Parafilm as a skin-substitute to cover a small amount of warmed animal blood [17]. This allows us to deliver DEET and quinine specifically either on the surface of the Parafilm or in the blood, an experiment not feasible to conduct with live human subjects. In this assay, we observed that both DEET and quinine were effective anti-feedants when mixed into blood, but only DEET completely blocked feeding when applied to the surface of the Glytube (Fig. 1J). These results agree with recent findings from *Culex quinquefasciatus* mosquitoes, which demonstrated that animals spent less time feeding on Parafilm-covered blood-soaked cotton balls if DEET was mixed into the blood [11]. These data support the hypothesis that contact DEET repellency is independent of bitter taste, and that an unknown, sensory mechanism repels *Ae. aegypti* mosquitoes on contact.

### The tarsi, not the proboscis, are required for contact DEET repellency

In search of the sensory appendages responsible for DEET contact repellency, we focused on the proboscis and legs because they are the primary appendages that contact the skin during landing (Fig. 1G). To test if the proboscis is sufficient to mediate contact DEET repellency, we modified the arm-in-cage assay and restricted the area of skin available for the mosquitoes to contact (Fig. 2A-B). The ~1.5 mm diameter circle of exposed skin we used in this assay is smaller than the distance between a mosquito’s forelegs, and she therefore cannot touch the skin with both her proboscis and her legs at the same time (Fig. 2B-C). In this assay, *orco* mutant mosquitoes blood-fed equally on solvent- and DEET-treated arms, suggesting that they are unable to sense DEET if only the proboscis touches the skin (Fig. 2C). In contrast, if both the legs and the proboscis can contact the skin (Fig. 2D), DEET remained an effective contact repellent (Fig. 2E). These data provide evidence that the proboscis is not sufficient to deter mosquitoes from biting DEET-treated human skin.

We next investigated if the legs are required to sense DEET on skin. Only the terminal segments of the leg, the tarsi, contact the skin (Fig. 2F). Tarsi are covered in sensory hairs called sensilla, which have pores that allow tastants to enter and activate sensory neurons [18] (Fig. 2G-I). We carried out experiments that asked whether some or all legs mediate DEET contact repellency. Initial experiments to surgically remove all tarsi were uninterpretable because the tarsi are required to produce the necessary force and leverage to pierce the skin [19]. To disrupt tarsal chemosensation without removing the tarsi, we coated them with UV-curing glues which have been used previously to occlude sensilla in taste organs [20] and antennae [21] in *D. melanogaster* flies (Fig. 2H-I).

When all tarsi were occluded by gluing, mosquitoes were no longer repelled by DEET-treated skin and bit DEET- and solvent-treated arms equally (Fig. 2J). Animals that were sham-treated or with their tibia glued were still repelled by DEET on contact (Fig. 2J), suggesting that the tarsi are necessary for contact DEET repellency. While observing the animals interact with these small areas of available skin surface, we noticed that they did not always contact the skin with tarsi on all 6 legs (Fig. 2D, 2G). We therefore asked if any pair of tarsi was dispensable or required for contact DEET repellency. Leaving any pair of tarsi unoccluded was sufficient to decrease biting events (Fig. 2K), suggesting that any pair of tarsi is sufficient to deter blood feeding on DEET-treated arms. This further suggests that the chemosensory neurons and receptors that sense DEET must be present in tarsi on all six legs.

**Figure 2.**
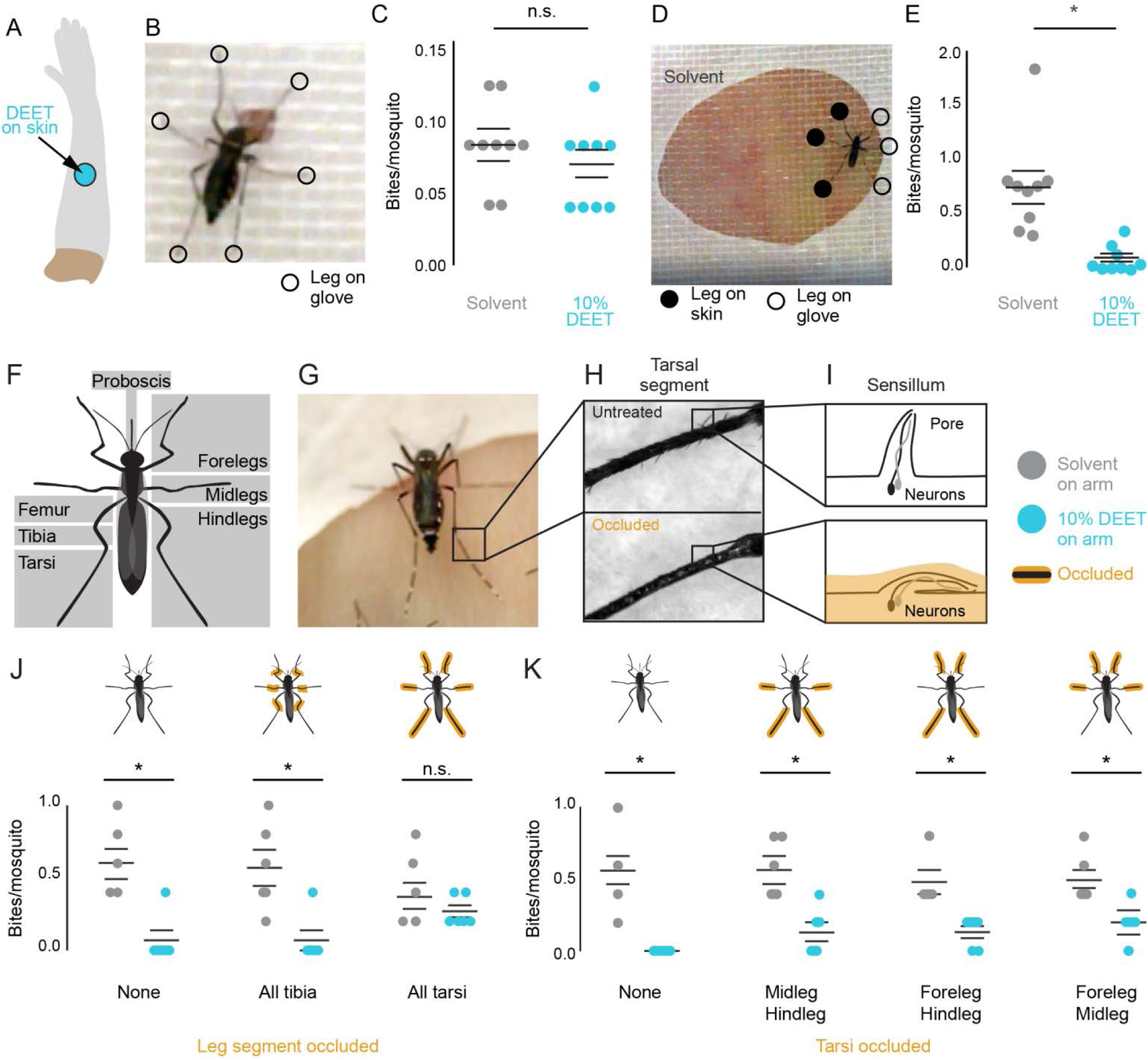
| The tarsi, not the proboscis, are required for contact DEET repellency. (A) Schematic of a DEET-treated arm with a 25 mm circle of accessible skin. (B) Video still of a mosquito feeding on a DEET-treated arm with a 1.5 mm circle of accessible skin. (C) Average number of biting events per mosquito on solvent- (gray) or DEET- (blue) treated skin (N=9 assays, 23-25 females/assay). (D) Video still of a mosquito feeding on a solvent-treated arm through a 25 mm circle of accessible skin, with positions of legs manually scored. (E) Mosquito biting events on solvent- (gray) or DEET- (blue) treated skin (N=9 assays, 23-25 females/assay). (F) Schematic of mosquito leg anatomy and the proboscis. (G) Video still of a mosquito on a human arm, highlighting the tarsi. (H) Examples of untreated (top) or UV glue occluded (bottom) tarsal segments. (I) Schematic of tarsal sensillum after occlusion by UV glue. (J-K) Mosquito biting events on solvent- (gray) or DEET- (blue) treated arms. Cartoons at the top indicate which appendages were occluded (N=6 assays, 5 females/assay). In C, E, J, and K horizontal lines represent mean ± SEM. Statistical significance was assessed with Student’s t-test (^∗^ p<0.05; n.s., not significant).

## Conclusions

DEET is a small, synthetic molecule that is the world’s most effective and widely used insect repellent [22]. Developed in World War II to protect soldiers threatened by mosquito-borne diseases such as malaria and yellow fever, DEET has been in civilian use for over 70 years [23]. It protects humans against bites from animals across vast evolutionary distances, including land leeches [24], ticks [25], and mosquitoes [26, 27]. Although highly effective, it has several undesirable properties that limit its use in areas of active mosquito-borne illnesses: it is oily on the skin and must be reapplied liberally at very high concentrations on all areas of exposed skin every 6 hours. This is impractical in the tropical zones where pathogen-infected mosquitoes are most dangerous. Despite various efforts to improve upon DEET, it remains the gold standard for personal protection. Remarkably, its mechanism of action is still incompletely understood, and this gap in our knowledge prevents the rational design of new highly effective molecules that address the deficiencies of DEET.

Our work challenges the previously accepted hypothesis that DEET acts solely as a disruptor of olfaction and as a bitter tastant, and discover a third mechanism by which mosquitoes sense DEET. We describe a mechanism of contact chemorepellency mediated by the tarsal segments of the leg, and show that this is the true sensory basis of repellency on skin. Further, we show that this tarsal contact repellency, not bitter taste sensed via the proboscis upon ingestion, is the relevant pathway used by mosquitoes to avoid DEET on skin. This is important because ongoing efforts to improve upon DEET as a multi-modal repellent molecule are based on the incorrect assumption that the contact effects of DEET act via an ingestive gustatory bitter tastant mechanism. Further investigation of the receptors and sensory neurons on the tarsi, not proboscis, are key to the understanding of the effectiveness of DEET and may aid in the development of new insect repellents. Moreover, it is a cautionary tale against using results from a non-biting insect like *D. melanogaster* flies to draw conclusions about mosquito behavior. Flies do not bite humans, and mosquitoes have evolved specialized sensing mechanisms relevant to their lifestyle as blood-feeding insects.

## Acknowledgments

We thank Josie Clowney, Matthew DeGennaro, Itzel Ishida, Xin Jin, Kevin Lee, and members of the Vosshall lab for discussion and comments on the manuscript, Matthew DeGennaro for early discussions, Vineeta Reddy for technical assistance with experiments in Figure 1B-D, and Tim Dennis for blinding all videos before analysis. Research reported in this publication was supported by the National Institute on Deafness and Other Communication Disorders of the National Institutes of Health under Award Number F31DC014222 awarded to E.J.D. Supported in part by grant # UL1 TR000043 from the National Center for Advancing Translational Sciences (NCATS, National Institutes of Health (NIH) Clinical and Translational Science Award (CTSA) program. L.B.V. is an investigator of the Howard Hughes Medical Institute.

## Author Contributions

Supervision, L.B.V; Investigation, E.J.D; Conceptualization, Methodology, Writing, and Funding Acquisition E.J.D. and L.B.V.

## Declaration of Interests

The authors declare no competing interests.

## STAR Methods

### CONTACT FOR REAGENT AND RESOURCE SHARING

Further information and requests for resources and reagents should be directed to and will be fulfilled by the Lead Contact, Leslie Vosshall.

### EXPERIMENTAL MODEL AND SUBJECT DETAILS

#### Mosquito rearing and maintenance

*Aedes aegypti orco^5/16^* heteroallelic mutants were generated by crossing homozygous *orco^5/5^* and *orco^16/16^* mutants and collecting F1 progeny [6]. Mosquitoes were reared at 25-28°C, 70%-80% relative humidity with a photoperiod of 14 h light: 10 h dark (lights on 7 AM) as previously described [6]. Eggs were hatched in deoxygenated, deionized water containing powdered Tetramin tablets (fish food) (Tetra; 16110M, Pet Mountain). Larvae were fed additional Tetramin until pupation. Pupae were placed in a small cup of deionized water, moved to a 30 cm^3^ cage (211261, Bugdorm), and allowed to eclose. Adult mosquitoes were housed with siblings with unlimited access to 10% sucrose (57-50-1, Thermo Fisher) (weight:volume in deionized water), and blood-fed on mice for stock maintenance. Blood-feeding on mice was approved and monitored by The Rockefeller University Institutional Animal Care and Use Committee (protocol 15772). Human subjects provided written, informed consent to participate in experiments, and these procedures were approved and monitored by The Rockefeller University Institutional Review Board (protocol LV-0652).

### METHOD DETAILS

#### Mosquito behavior experiments

All behavioral experiments were carried out with 7-14 day old *orco^5/16^* heteroallelic mutant female mosquitoes that had not previously taken a blood-meal. All assays were carried out at Zeitgeber time (ZT) 6-ZT10 at 25-28°C and 70-80% humidity. Unless otherwise stated, cold anesthesia was carried out by working with animals in a 4° C cold room. Blood-feeding was scored by eye by identifying large red-pigmented abdomens. No partially blood-fed mosquitoes were observed in any assays where end-of-assay scoring was used.

#### CAFE feeding assay

Animals were sexed and sorted under cold anesthesia (4° C) and fasted for 18-20 hours with access to water. This assay was adapted for the mosquito from similar assays for *Drosophila melanogaster* [13], and as described previously [12]. At the start of each trial, five fasted female mosquitoes were transferred by mouth pipette to a polypropylene vial (89092–742, VWR) with access to two 5 mL calibrated glass capillaries (53432–706, VWR) embedded in cotton plugs (49-101, Genesee Scientific) and barely protruding from the bottom of the plug surface. A small piece of red tape (89097-932, VWR) was affixed to the bottom of the plug, because previous work demonstrated that this increased participation in the assay [12]. One capillary served as the control, containing 10% sucrose (weight:volume) in deionized water supplemented with 1% ethanol solvent (E7023, Millipore Sigma). The stimulus capillary contained 10% sucrose supplemented with one of the following chemicals: 1% DEET (CID 24893319; D100951, Sigma-Aldrich), 1 mM lobeline (CID 101615; 141879, Millipore Sigma), or 5 mM quinine (CID 16211610, Q1250, Millipore Sigma). These were prepared from 100X stock solutions in ethanol for bitter tastants, or 50% DEET, such that the final concentration of
ethanol in 10% sucrose was 1%. After four hours, the remaining liquid in all capillaries was measured by a blinded observer in millimeters, by aligning a metric ruler to the tip of the capillary and measuring the height of the liquid meniscus. Mosquito-less vials with 2 capillaries filled with 10% sucrose and 1% ethanol served as evaporation controls. Eight evaporation vials were used each day. An average evaporation amount for each day of experiments was calculated (EVAP) by first calculating an average of the two control capillaries’ evaporation for each vial, and then averaging across all evaporation control vials on that day. For each test vial, the reduction in liquid level was recorded for the 10% sucrose capillary (CONTROL) and the 10% sucrose + stimulus capillary (STIMULUS). The preference index was calculated as follows: [(STIMULUS – EVAP) – (CONTROL – EVAP)] / [(STIMULUS – EVAP) + (CONTROL – EVAP)]. Vials were blinded before manual scoring. Vials were excluded if any of the 5 animals died during the assay. No artificial CO_2_ was added to these experiments.

#### Glytube blood-feeding assay

Animals were sexed and sorted under cold anesthesia (4° C) into groups of 15-16 females and fasted for 18-24 hours with access to water. Mosquitoes were provided defibrinated sheep blood (DSB500, Hemostat Laboratories) warmed in a 42° C water bath using Glytube membrane feeders as described [17]. No synthetic CO_2_ was added to these cages but assays were carried out in close proximity to a breathing human. DEET or quinine were either applied to the surface by dipping the Glytube into a DEET or quinine solution, or mixed into the blood immediately prior to the start of the assay. Glytubes were placed directly on the mesh tops of the cups housing each group of mosquitoes so that they were flush with the surface of the mesh, easily accessible for the mosquitoes to touch and puncture for feeding. Animals were allowed to feed for 15 minutes, and then the Glytubes were removed, and animals scored for blood-feeding status. These assays were not blinded.

#### Human blood-feeding and biting assays

Standard arm-in-cage biting assays [15, 16] were carried out with modifications as previously described [6] and additional modifications in each section below.

The arm-in-cage blood feeding assays were not filmed, because they were endpoint assays in which blood-feeding was scored. All other assays described in this section used a Canon EOS60D camera at 60 fps directed 90 degrees from the arm. The camera lens was inserted into a cage through a mesh sleeve opening. An arm was either pressed against the opposing side of the cage (constrained feeding access assay) or inserted into the cage through a mesh sleeve opening on the wall adjacent to the camera. The arm was positioned in front of the camera, through the middle of the cage, as drawn in Fig. 1E.

Videos were blinded in groups of eight or twelve by a volunteer who changed the names of the videos. The blinded videos were then scored, frame by frame, to record the number of skin contacts and bites. A contact was defined as when a mosquito landed on the skin, observed by her contacting the skin with at least one tarsi or proboscis while her wings had stopped moving. If no contacts were observed, the video was discarded. This was rare (<2% of videos). A bite was defined subjectively but required that: (1) the proboscis was in contact with the skin (2) the mosquito forelegs and midlegs were stationary (3) the mosquito head did not move, and then a slight sawing motion was visible. Whenever possible, the bite was confirmed by noting that the skin reddened, although some bites occurring at the very end of an assay could not be confirmed in this way. Bite scoring did not require visual detection of blood in the abdomen, as this was often difficult either because of the duration of the feeding event (a bite at 9:45 of a 10:00 video) and the body positioning of the animal.

#### Arm-in-cage blood-feeding assays

To present DEET or bitters to mosquitoes on a live human forearm, a 25 mm diameter hole either at the base of the wrist or 75 mm above the wrist, closer to the elbow on the inside of the forearm, was cut into an elbow-length latex glove (19-668-001, Fisher Scientific). To prepare the arm, three horizontal lines were drawn approximately 48, 50, and 52 mm above the wrist in ethanol-soluble ballpoint pen ink. 0.5 mL of either solvent or a test substance in solvent (lobeline, quinine, or DEET) was added to the upper or lower forearm of a human volunteer (27-year-old female) before donning the glove. The test substance was applied such that the closest ballpoint pen ink line was smeared, while the middle line was not. If the lower half of the forearm was treated, the lowest ballpoint pen ink line was smeared and no others. A glove with a hole at the base of the wrist was used, exposing a 25 mm diameter area of solvent- or test-treated skin.

A group of 25 females was released into a 30 cm^3^ Bugdorm cage (Bugdorm), and given five minutes to acclimate to the cage. The gloved arm was then placed in the cage for ten minutes. After ten minutes, the arm was removed and cage moved to a 4° C cold room to anesthetize the animals. Animals were scored as blood-fed or non-blood-fed based by visual inspection of the abdomen. No synthetic CO_2_ was added to these cages but assays were carried out in close proximity to a breathing human. These assays were blinded by re-labeling the test substances before application to the arm. These assays were pseudo-randomized such that the stimuli were provided in a different order each day, on different arm halves, and a solvent control was included each day of experimentation. The per cent blood-fed was calculated by counting the number of fed mosquitoes divided by the total number of mosquitoes, multiplied by 100.

#### Constrained feeding access assay

This assay is a modification of the arm-in-cage assay, where the gloved human arm exposing either 25 mm or 1.5 mm of skin is instead pressed against the mesh on the outside of the cage. By decreasing the surface area that the mosquitoes could explore before finding the hole in the glove, participation in the small hole (1.5 mm) trials was increased, and DEET remained effective in the large hole (25 mm) trials, demonstrating that the mosquitoes were able to bite through the mesh easily. These assays were not blinded and were pseudo-randomized such that the stimuli were provided in a different order each day, and a solvent control was included each day of experimentation. The videos were blinded before manual annotation by re-naming the files. Bites/mosquito was calculated for each video by dividing the number of animals biting by the number of animals in the assay.

#### Arm-in-cage mosquito leg occlusion biting assays

Animals were sexed and sorted into cups in groups of five using a mouth aspirator, and anesthetized by placing the cups on wet ice. Individual mosquitoes were transferred to a petri dish filled with wet ice. Tarsi were glue-occluded by inserting them one at a time into the narrow end of a 1 mL pipette tip containing 200-500 μL UV curing glue (KOA 300-1, Kemxert), coating the legs, which were then removed from the pipette tip and cured with a 405 nm 5 mW laser pointer (QQ-Tech) for 20 seconds with the laser pointer held approximately 25 mm away from the tip of the tarsi, and pointed toward the abdomen to illuminate the whole tarsi.

Unoccluded controls were handled identically with the exception that the pipette tip was empty, so no glue was applied. Tibia were glued by slowly applying UV glue with a 200 μL pipette tip until coated, then cured for 20 seconds as described above. The process for each animal took 2-5 minutes. Animals with the same treatment were housed in groups of 5 females for 18-24 hours with access to water at 25-28°C and 70-80% humidity. If any animals died overnight, that group of females was discarded. This was a rare occurrence. Bites/mosquito was calculated for each video by dividing the number of animals biting by the number of animals in the assay.

## QUANTIFICATION AND STATISTICAL ANALYSIS

R version 3.3.2 Sincere Pumpkin Patch (CRAN) was used for all statistical analyses. Statistical details including exact values of N and what N represents are indicated in the figure legends and any calculations are defined in the method details. Significance was defined as p<0.05. Sample sizes were estimated by a power analysis on pilot data, with the exception of occlusion experiments and Glytube feeding experiments, which were based on sample sizes of previous studies. Exclusion criteria, blinding, and randomization for each behavioral assay are defined in the method details.

## DATA AND SOFTWARE AVAILABILITY

Software and custom scripts used for statistical analysis, plotting, and manual video annotation are listed in the Key Resources Table. All data in the paper are available in Supplemental Data File 1, with the exception of raw video files, which are available upon request.

KEY RESOURCES TABLE

KEY RESOURCES TABLE
| REAGENT or RESOURCE | SOURCE | IDENTIFIER |
| --- | --- | --- |
| Biological Samples |  |  |
| Defibrinated sheep blood | Hemostat Laboratories | DSB500 |
| Chemicals |  |  |
| DEET (N,N diethyl-meta-toluamide) CID: 4284 | Millipore Sigma | D100951 |
| Lobeline CID: 101615 | Millipore Sigma | 141879 |
| Quinine CID: 16211610 | Millipore Sigma | Q1250 |
| Ethanol (solvent) CID: 329799002 | Millipore Sigma | E7023 |
| Sucrose CID: 5988 | Thermo Fisher | CAS 57-50-1 |
| ATP CID: 5957 | Millipore Sigma | A3377-GG |
| Deposited Data |  |  |
| Data and analyses | This Paper | Supplemental Data File 1 |
| Experimental Models: Organisms |  |  |
| *Aedes aegypti orco5* mutant mosquitoes | BEI Resources | NR-44377 |
| *Aedes aegypti orco16* mutant mosquitoes | BEI Resources | NR-44378 |
| Software and Algorithms |  |  |
| R | CRAN | https://www.r-project.org/ |
| plyr (Rpackage) | CRAN | RRID:SCR_001905 |
| tidyverse (Rpackage) | CRAN | https://cran.r-project.org/web/packages/tidyverse/index.html |
| multcompView (Rpackage) | CRAN | https://CRAN.R-project.org/package=multcompView |
| ggthemes (Rpackage) | CRAN | https://CRAN.R-project.org/package=ggthemes |
| reshape2 (Rpackage) | CRAN | http://www.jstatsoft.org/v21/i12/ |
| Other |  |  |
| 5 μL glass capillaries | VWR | 53432-706 |
| Cotton flugs | Genessee Scientific | 49-102 |
| Narrow plastic vials | Genessee Scientific | 32-116 |
| Cups | Webstaurant | KH16A-J8000 |
| Mesh | McMaster Carr | 98315K58 |
| Tetramin fish food | Pet Mountain | 16110M |
| 200 μL pipette tips, with filter | Fisher Scientific | 21-377-354 |
| 1 mL pipette tips, no filter | Sigma Aldrich | CLS4868 |
| Time tape (red) | VWR | 89097-932 |
| 405 nm 5 mW laser pointer | QQ-Tech | Discontinued |
| KOA 300 UV curing glue | Kemxert | KOA 300-1 |
| Canon EOS60D | Canon | EOS60D |
| 30 cm<sup>3</sup> BugDorm Insect Rearing Cage | Bugdorm | 211261 |
| BioClean Nerva Extra Length Nitrile Cleanroom Gloves | Thermo Fisher | 19-668-001 |

## Supplemental Information

All raw data in this paper are available in Supplemental Data File 1.

---

**Graphical Abstract**

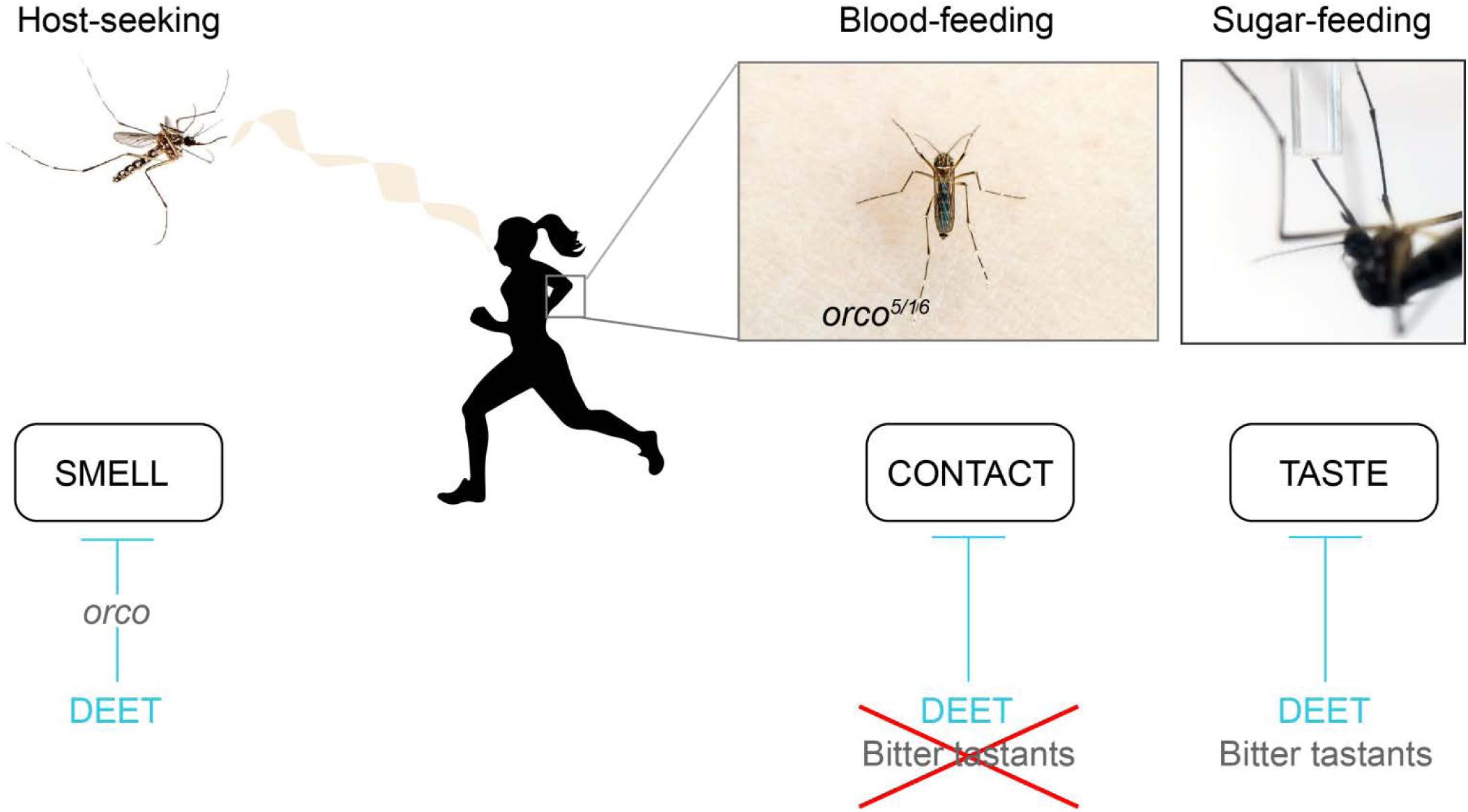

## References

1. DeGennaro, M. (2015). The mysterious multi-modal repellency of DEET. Fly (Austin) 9, 45–51.

2. Ditzen, M., Pellegrino, M., and Vosshall, L.B. (2008). Insect odorant receptors are molecular targets of the insect repellent DEET. Science 319, 1838–1842.

3. Liu, C., Pitts, R.J., Bohbot, J.D., Jones, P.L., Wang, G., and Zwiebel, L.J. (2010). Distinct olfactory signaling mechanisms in the malaria vector mosquito *Anopheles gambiae*. PLoS Biol 8.

4. Pellegrino, M., Steinbach, N., Stensmyr, M.C., Hansson, B.S., and Vosshall, L.B. (2011). A natural polymorphism alters odour and DEET sensitivity in an insect odorant receptor. Nature 478, 511–514.

5. Syed, Z., and Leal, W.S. (2008). Mosquitoes smell and avoid the insect repellent DEET. Proc Natl Acad Sci U S A 105, 13598–13603.

6. DeGennaro, M., McBride, C.S., Seeholzer, L., Nakagawa, T., Dennis, E.J., Goldman, C., Jasinskiene, N., James, A.A., and Vosshall, L.B. (2013). *orco* mutant mosquitoes lose strong preference for humans and are not repelled by volatile DEET. Nature 498, 487–491.

7. Abramson, C.I., Giray, T., Mixson, T.A., Nolf, S.L., Wells, H., Kence, A., and Kence, M. (2010). Proboscis conditioning experiments with honeybees, *Apis mellifera caucasica*, with butyric acid and DEET mixture as conditioned and unconditioned stimuli. J Insect Sci 10, 122.

8. Lee, Y., Kim, S.H., and Montell, C. (2010). Avoiding DEET through insect gustatory receptors. Neuron 67, 555–561.

9. Klun, J.A., Khrimian, A., and Debboun, M. (2006). Repellent and deterrent effects of SS220, Picaridin, and Deet suppress human blood feeding by *Aedes aegypti*, *Anopheles stephensi*, and *Phlebotomus papatasi*. J Med Entomol 43, 34–39.

10. Sanford, J.L., Shields, V.D.C., and Dickens, J.C. (2013). Gustatory receptor neuron responds to DEET and other insect repellents in the yellow-fever mosquito, *Aedes aegypti*. Naturwissenschaften 100, 269–273.

11. Lu, W., Hawang, J.K., Zeng, F., and Leal, W.S. (2017). DEET as a feeding deterrent. PLoS One 12, e0189243.

12. Corfas, R.A., and Vosshall, L.B. (2015). The cation channel TRPA1 tunes mosquito thermotaxis to host temperatures. Elife 4, e11750.

13. Ja, W.W., Carvalho, G.B., Mak, E.M., de la Rosa, N.N., Fang, A.Y., Liong, J.C., Brummel, T., and Benzer, S. (2007). Prandiology of *Drosophila* and the CAFE assay. Proc Natl Acad Sci U S A 104, 8253–8256.

14. Liesch, J., Bellani, L.L., and Vosshall, L.B. (2013). Functional and genetic characterization of neuropeptide Y-like receptors in *Aedes aegypti*. PLoS Negl Trop Dis 7, e2486.

15. Schreck, C.E. (1977). Techniques for the evaluation of insect repellents: a critical review. Annu Rev Entomol 22, 101–119.

16. Logan, J.G., Stanczyk, N.M., Hassanali, A., Kemei, J., Santana, A.E., Ribeiro, K.A., Pickett, J.A., and Mordue Luntz, A.J. (2010). Arm-in-cage testing of natural human-derived mosquito repellents. Malar J 9, 239.

17. Costa-da-Silva, A.L., Navarrete, F.R., Salvador, F.S., Karina-Costa, M., Ioshino, R.S., Azevedo, D.S., Rocha, D.R., Romano, C.M., and Capurro, M.L. (2013). Glytube: a conical tube and parafilm M-based method as a simplified device to artificially blood-feed the dengue vector mosquito, *Aedes aegypti*. PLoS One 8, e53816.

18. McIver, S., and Siemicki, R. (1978). Fine structure of tarsal sensilla of *Aedes aegypti* (L.) (Diptera: Culicidae). J. Morphol 155, 137–156.

19. Jones, J.C., and Pilitt, D.R. (1973). Blood-feeding behavior of adult *Aedes aegypti* mosquitoes. Biol. Bull. 145, 127–139.

20. Olsen, S.R., and Wilson, R.I. (2008). Lateral presynaptic inhibition mediates gain control in an olfactory circuit. Nature 452, 956–960.

21. Wasserman, S., Salomon, A., and Frye, M.A. (2013). *Drosophila* tracks carbon dioxide in flight. Curr Biol 23, 301–306.

22. Katz, T.M., Miller, J.H., and Hebert, A.A. (2008). Insect repellents: historical perspectives and new developments. J Am Acad Dermatol 58, 865–871.

23. McCabe, E.T., Barthel, W.F., Gertler, S.I., and Hall, S.A. (1954). Insect Repellents. III. N, N-diethylamides. J. Org. Chem. 19, 493–498.

24. Tawatsin, A., Thavara, U., Chansang, U., Chavalittumrong, P., Boonruad, T., Wongsinkongman, P., Bansidhi, J., and Mulla, M.S. (2006). Field evaluation of deet, Repel Care, and three plant based essential oil repellents against mosquitoes, black flies (Diptera: Simuliidae) and land leeches (Arhynchobdellida: Haemadipsidae) in Thailand. J Am Mosq Control Assoc 22, 306–313.

25. Evans, S.R., Korch, G.W., Jr., and Lawson, M.A. (1990). Comparative field evaluation of permethrin and deet-treated military uniforms for personal protection against ticks (Acari). J Med Entomol 27, 829–834.

26. Bernier, U.R., Furman, K.D., Kline, D.L., Allan, S.A., and Barnard, D.R. (2005). Comparison of contact and spatial repellency of catnip oil and N,N-diethyl-3-methylbenzamide (deet) against mosquitoes. J Med Entomol 42, 306–311.

27. Thavara, U., Tawatsin, A., Chompoosri, J., Suwonkerd, W., Chansang, U.R., and Asavadachanukorn, P. (2001). Laboratory and field evaluations of the insect repellent 3535 (ethyl butylacetylaminopropionate) and deet against mosquito vectors in Thailand. J Am Mosq Control Assoc 17, 190–195.

